# *F*_ST_ and kinship for arbitrary population structures I: Generalized definitions

**DOI:** 10.1101/083915

**Authors:** Alejandro Ochoa, John D. Storey

## Abstract

*F*_ST_ is a fundamental measure of genetic differentiation and population structure, currently defined for subdivided populations. *F*_ST_ in practice typically assumes *independent, non-overlapping subpopulations*, which all split simultaneously from their last common ancestral population so that genetic drift in each subpopulation is probabilistically independent of the other subpopulations. We introduce a generalized *F*_ST_ definition for arbitrary population structures, where individuals may be related in arbitrary ways, allowing for arbitrary probabilistic dependence among individuals. Our definitions are built on identity-by-descent (IBD) probabilities that relate individuals through inbreeding and kinship coefficients. We generalize *F*_ST_ as the mean inbreeding coefficient of the individuals’ local populations relative to their last common ancestral population. We show that the generalized definition agrees with Wright’s original and the independent subpopulation definitions as special cases. We define a novel coancestry model based on “individual-specific allele frequencies” and prove that its parameters correspond to probabilistic kinship coefficients. Lastly, we extend the Pritchard-Stephens-Donnelly admixture model in the context of our coancestry model and calculate its *F*_ST_. To motivate this work, we include a summary of analyses we have carried out in follow-up papers, where our new approach has been applied to simulations and global human data, showcasing the complexity of human population structure, demonstrating our success in estimating kinship and *F*_ST_, and the shortcomings of existing approaches. The probabilistic framework we introduce here provides a theoretical foundation that extends *F*_ST_ in terms of inbreeding and kinship coefficients to arbitrary population structures, paving the way for new estimators and novel analyses.

Note: This article is Part I of two-part manuscripts. We refer to these in the text as Part I and Part II, respectively.

**Part I:** Alejandro Ochoa and John D. Storey. “*F*_ST_ and kinship for arbitrary population structures I: Generalized definitions”. *bioRxiv* (10.1101/083915) (2019). https://doi.org/10.1101/083915. First published 2016-10-27.

**Part II:** Alejandro Ochoa and John D. Storey. “*F*_ST_ and kinship for arbitrary population structures II: Method of moments estimators”. *bioRxiv* (10.1101/083923) (2019). https://doi.org/10.1101/083923. First published 2016-10-27.

## 1 Introduction

A population of mating organisms is *structured* if its individuals do not mate randomly, which results in an increase in mean homozygozity over the population compared to that of a randomly mating population [3, 4]. *F*_ST_ is a parameter that measures population structure [5, 6], which is typically understood through homozygosity. An unstructured population has *F*_ST_ = 0 and genotypes at each locus have Hardy-Weinberg proportions. At the other extreme, a fully differentiated population has *F*_ST_ = 1 and every subpopulation at every locus is homozygous for some allele. In addition to measuring population differentiation, *F*_ST_ is also used to model DNA profile matching uncertainty in forensics [7–13] and to identify loci under selection [14–21]. Current *F*_ST_ definitions assume a partitioned or subdivided population into discrete, non-overlapping subpopulations [5, 6, 22–24]. Many *F*_ST_ estimators further assume that subpopulations have evolved independently from the most recent common ancestor (MRCA) population [21–24], which occurs only if every subpopulation split from the MRCA population at the same time (Fig. 1A, Fig. 2A). However, populations such as humans are not naturally subdivided [11, 25–27] (Fig. 1B); thus, arbitrarily imposed subdivisions may yield correlated subpopulations that no longer satisfy the independent subpopulations model assumed by existing *F*_ST_ estimators (Fig. 2B). In this work, we build a generalized *F*_ST_ definition applicable to arbitrary population structures, including arbitrary evolutionary dependencies.

**Figure 1:**
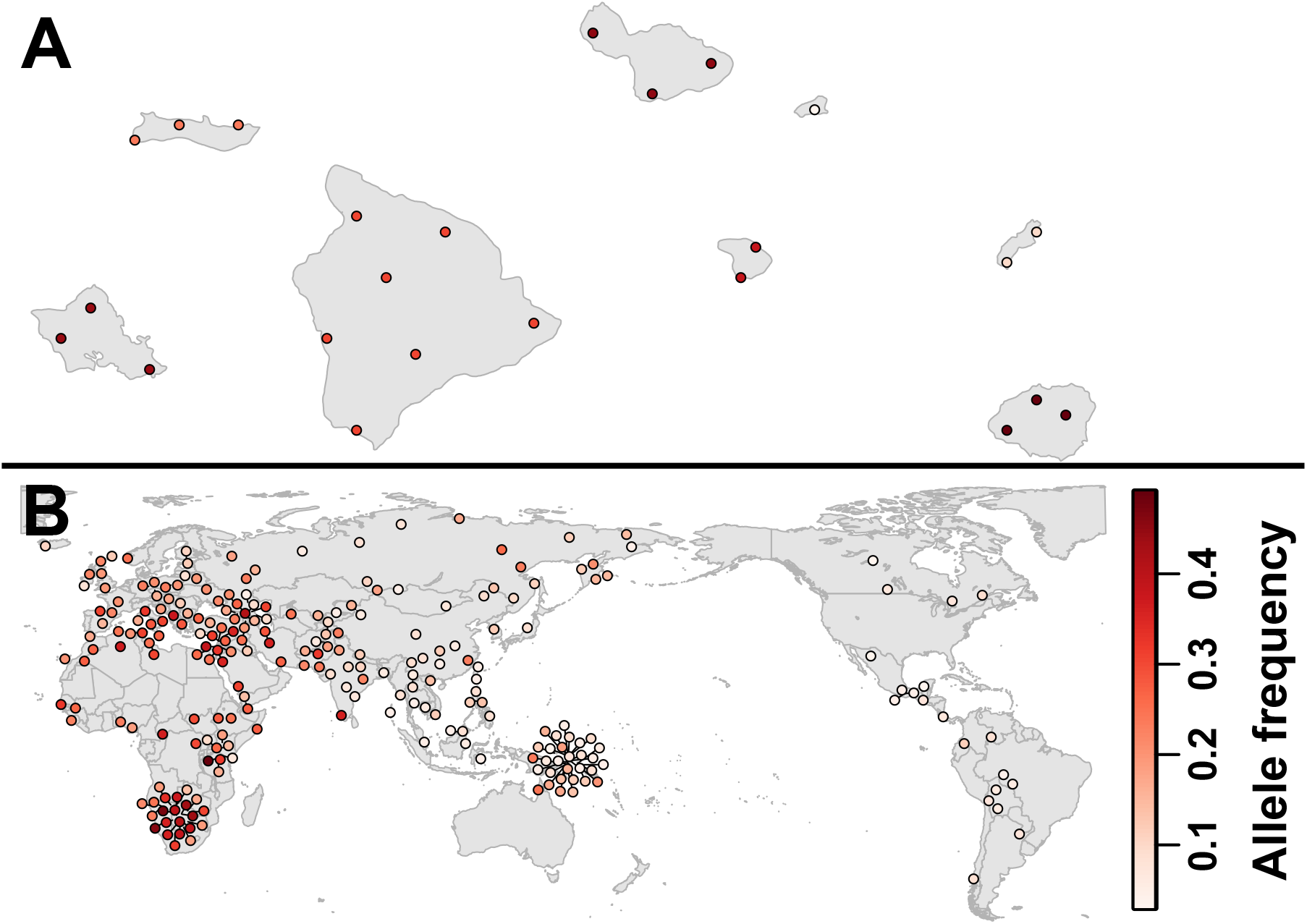
Independent subpopulations model versus median-*F*_ST_ human SNP. In these maps, circles are subpopulations (moved to prevent overlaps in panel B) colored by their mean allele frequency (AF) at a locus. **A.** A simulated locus from the independent subpopulations model illustrated using islands. Individuals from the same island draw their alleles from the same pool, so they have the same underlying AF, while individuals from different islands evolve independently (AFs across islands are uncorrelated). **B.** AFs at SNP locus rs2650044 in the Human Origins datasets of [28–30], illustrates typical differentiation in humans. This locus had the median per-locus *F*_ST_ estimate (≈ 0.0961) among loci with minor allele frequency ≥ 10% using the estimator of [22] and the *K* = 244 subpopulations shown. Since AFs display strong geographical correlation, the human population does not fit the independent subpopulations model. Subpopulation AFs are estimated using Empirical Bayes (see Supplementary Information, Section S3).

**Figure 2:**
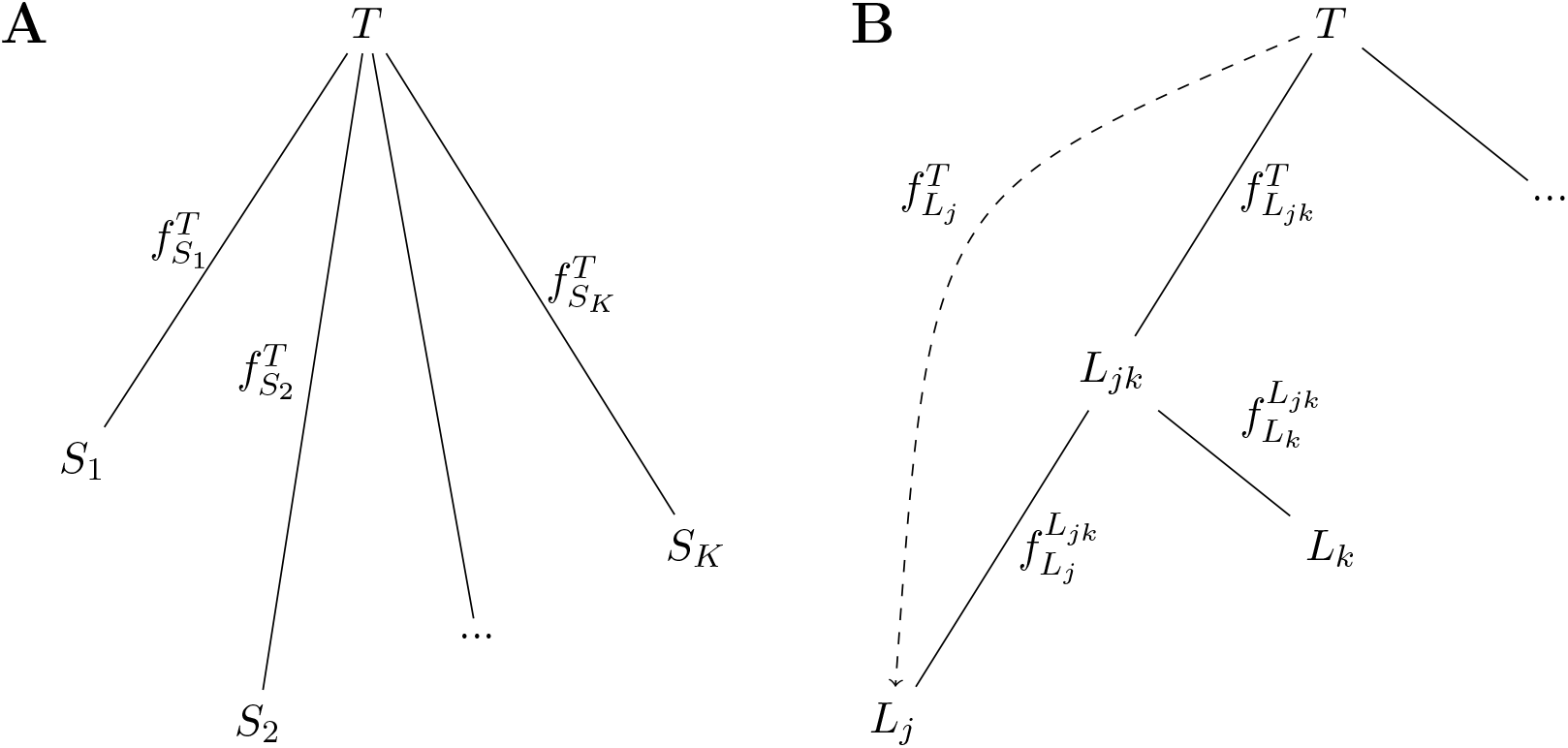
Independent subpopulations versus arbitrary population structures. These trees illustrate relationships (edges) between populations (nodes). The edge length between two populations *T* and *S* is proportional to the inbreeding coefficient 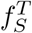 (see Section 3.1). **A.** In the independent subpopulations model, *K* subpopulations *S*_1_,…, *S*_*K*_ evolved independently from *T*, which requires that every *S*_*u*_ split from *T* at the same time. **B.** In an arbitrary population structure, each individual *j* has its own local population *L*_*j*_, and every pair of individuals (*j, k*) have a jointly local population *L*_*jk*_ from which *L*_*j*_ and *L*_*k*_ evolved (see Section 3.2). We do not assume a bifurcating tree process: the case for three or more individuals is not a tree (only two individuals *j* and *k* are shown). 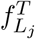 and 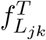 are relative to *T*, while 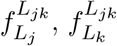 are relative to *L*_*jk*_.

Natural populations are often structured due to population size differences and the constraints of distance and geography [31]. For example, the genetic population structure of humans shows evidence of population bottlenecks migrating out of Africa [32–40] as well as numerous admixture events [41–45]. Notably, human populations display genetic similarity that decays smoothly with geographic distance, rather than taking on discrete values as would be expected for independent subpopulations [11, 27, 35, 37–39] (Fig. 1B). Current *F*_ST_ definitions do not apply to these complex population structures.

*F*_ST_ is known by many names, including fixation index [6] and coancestry coefficient [23, 46]). *F*_ST_ is also alternatively defined in terms of the variance of subpopulation allele frequencies [6], variance components [47], correlations [22], and genetic distance [46]. Our generalized *F*_ST_ is defined using inbreeding coefficients, like Wright’s *F*_ST_. There is also a diversity of summary statistics that measure locus-specific differentiation, such as *G*_ST_, 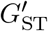, and *D*, which are functions of observed allele frequencies, and which approximate *F*_ST_ under certain conditions [48–53]. We consider *F*_ST_ as a genome-wide evolutionary parameter given by relatedness, which modulates the random drift of allele frequencies across loci but does not depend on these frequencies, mutation rates, or other locus-specific features. We review these previous *F*_ST_ definitions in greater detail in Supplementary Information, Section S1. The focus of our work is to generalize and accurately estimate the genome-wide *F*_ST_ in individuals with arbitrary relatedness, and does not presently concern locus-specific *F*_ST_ estimation or the identification of loci under selection.

The developments in this paper have lead to improved estimates of *F*_ST_ and kinship in Part II [2]. We have also applied these new probabilistic quantities and estimators to data from the Human Origins and 1000 Genomes Project data sets in ref. [54]. To motivate the generalized definitions we present in this work, in Section 2 we provide an overview of simulation results demonstrating the accuracy of the estimators (from Part II) and findings from analyzing the Human Origins and 1000 Genomes Project datasets (from ref [54]). These results establish that a generalized definition of *F*_ST_ in terms of kinship and inbreeding for arbitrary population structures is needed.

In Section 3 we formally define kinship and inbreeding coefficients, which measure how individuals are related, quantify population structure, and are the foundation of our work. We then generalize *F*_ST_ in terms of individual parameters (namely, inbreeding coefficients), and in analogy to Wright’s *F*_IS_, model local inbreeding on an individual basis. Our *F*_ST_ applies to arbitrary population structures, generalizing previous *F*_ST_ definitions restricted to subdivided populations.

In Section 4 we show a connection between the coalescent and kinship, inbreeding and generalized *F*_ST_. This provides a generalization of a previous result showing the relationship between the coalescent and the classic *F*_ST_ defined on subdivided populations. In Section 5 we define a coancestry model that parametrizes the correlations of “individual-specific allele frequencies” (IAFs), a recent tool that also accommodates individual-specific relationships [55, 56]. Our model is related to previous models between populations [23, 57]. We prove that our coancestry parameters correspond to kinship coefficients, thereby preserving their probabilistic interpretations, and we relate these parameters to *F*_ST_.

Lastly, in Section 6 we provide a novel *F*_ST_ analysis for admixed individuals by applying our coancestry model from Section 5 to the widely-used Pritchard-Stephens-Donnelly (PSD) admixture model, in which individuals derive their ancestry from several ancestral subpopulations with individual-specific admixture proportions [58–60]. We analyze an extension of the PSD model [55, 61–64] that generates allele frequencies from the Balding-Nichols distribution [7], and propose a more complete coancestry model for the ancestral subpopulations. We derive equations relating *F*_ST_ to the model parameters of PSD and its extensions. These results enable us to use an admixture simulation without independent subpopulations to benchmark kinship and *F*_ST_ estimators in Section 6 of Part II.

Our generalized definitions permit the analysis of *F*_ST_ and kinship estimators under arbitrary population structures, and pave the way forward to new estimation approaches, which are the focus of our following work in this series (Part II).

## 2 Motivating analyses

The results presented here lead to a deeper understanding of the limitations of existing *F*_ST_, kinship, and inbreeding estimators. Specifically, the assumptions underlying existing estimators are too restrictive and do not align with the properties of human populations that have been revealed through recent studies. In Part II, we theoretically calculate and then numerically verify complex biases that manifest from existing estimators when the population structure and relatedness violates the non-overlapping and independently evolving subpopulations assumptions. This then leads to new estimators of *F*_ST_, kinship, and inbreeding proposed in Part II. In ref. [54], we applied the estimators from Part II to data from the Human Origins study and 1000 Genomes Project (TGP). There, it is revealed on these seminal studies that the theory, methods, and simulations from Part I and Part II hold true on real data. Although the results summarized in this section involve details presented in full in Part II and ref. [54], it may be useful to the reader to see the ultimate consequences of the theory present in the current paper, Part I.

In Part II, we carried out simulations in two scenarios. The first scenario approximately satisfies the assumptions of the existing (Weir-Cockerham) estimate of *F*_ST_. The second scenario is an admixture model (described in Section 6), which reflects the characteristics we have observed in real data where there are no well-defined independent subpopulations. Fig. 3, columns A and B, show the results of these simulations. It can be seen that that both the existing and proposed estimators do well in the first scenario (Fig. 3A) where the population is divided into non-overlapping subpopulations that have independently evolved from a common ancestral population. However, in the second scenario (Fig. 3B) where these assumptions are violated, the existing estimators show notable downward bias. Our theoretical results determine exactly what this bias is for both kinship and *F*_ST_.

**Figure 3:**
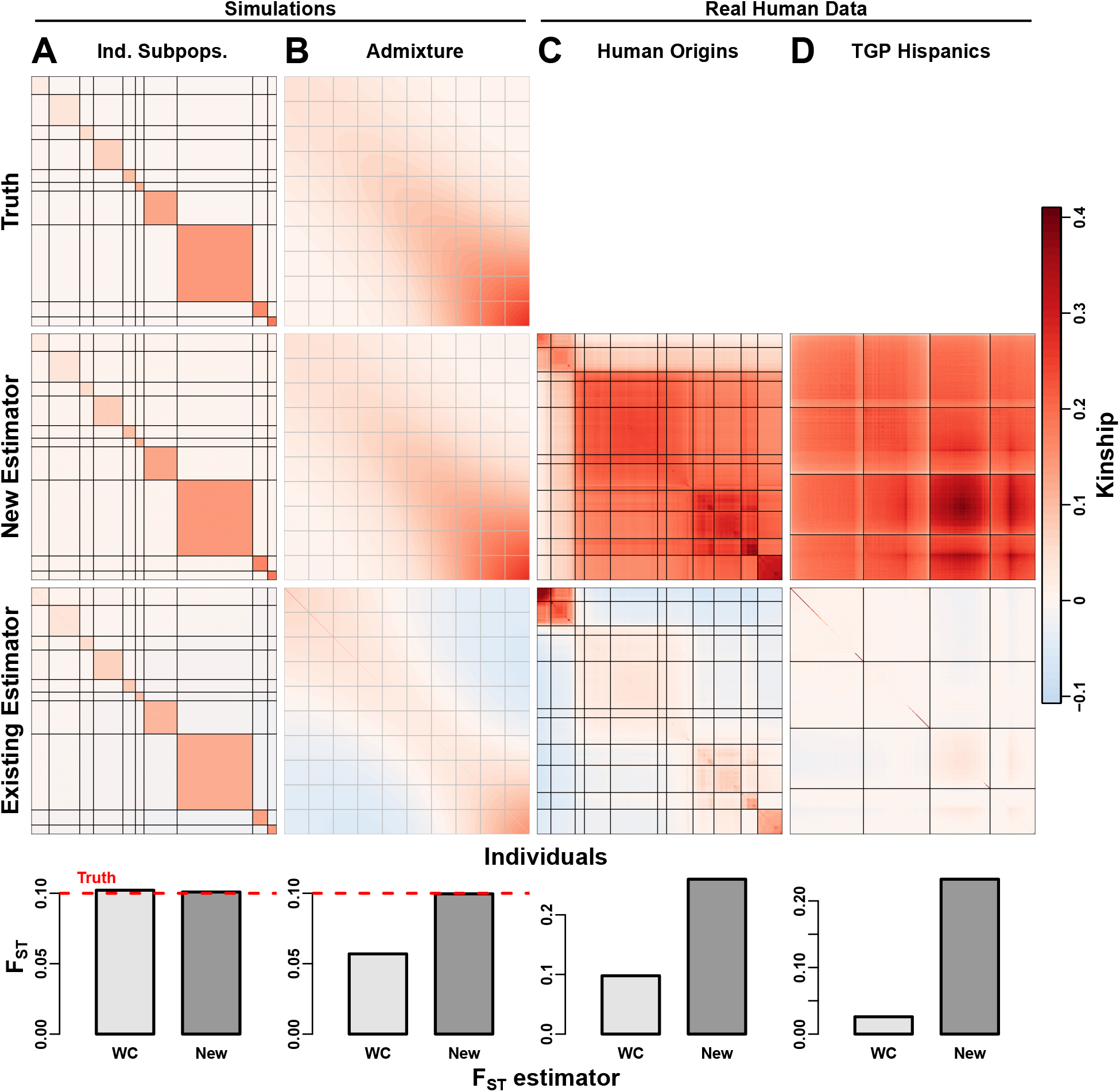
New and existing kinship and *F*_ST_ estimates in simulations and real human data. Each column corresponds to a given dataset and contains four panels: (1) the true kinship matrix (for simulations only; unknown in real data), (2) our new kinship estimates, (3) the standard kinship estimates, and (4) the comparison of the Weir-Cockerham (WC) *F*_ST_ estimates to our new *F*_ST_ estimates and the true *F*_ST_ value (red dashed lines; unknown in real data). Each kinship matrix plots the kinship values (color scale) between every pair of individuals (x and y axes) and the inbreeding values along the diagonal. **A.** The independent subpopulations simulation is the only scenario where existing *F*_ST_ estimators are unbiased. The standard kinship estimator has a small bias since the average kinship is fairly low. **B.** A spatial admixture simulation demonstrates biases in existing approaches (distortions in standard kinship estimates and *F*_ST_ estimates that are half of the true values) and superior performance of our kinship and *F*_ST_ approach (see Section 6 in Part II for simulation details). **C.** The Human Origins dataset for global human populations. Individuals were grouped into *K* = 11 continental clusters (see [54]). **D.** The Hispanic subset of the 1000 Genomes Project data. Individuals were grouped into *k* = 4 clusters by sampling location (see [54]).

In ref. [54], we then analyzed data from the Human Origins [28–30] and TGP studies [65], both of which consist of individuals sampled from a global distribution of ancestries. For the TGP data, we specifically limited our analysis to Hispanics. Our novel kinship estimates calculated on these data reveal a complex population structure in the global human population (Fig. 3C) and in Hispanics in particular (Fig. 3D). Since there are no independent subpopulations in the human data, existing kinship and *F*_ST_ estimates in these data will also be downwardly biased, which can be seen in the bottom two rows of Fig. 3C-D. In contrast, our more accurate novel *F*_ST_ estimates measure greater differentiation than has been previously reported (Fig. 3C-D, second and fourth rows). A deeper analysis of our calculations reveals a clear connection between our estimated kinship structure (but not existing estimates) and the global human migrations under the African Origins model [54]. Our results suggest that common population genetic analyses on real human data will greatly benefit from our improved kinship and *F*_ST_ estimation framework.

## 3 Generalized definitions in terms of individuals

Now that we have established the need for a more flexible population structure model that does not assume independent subpopulations, we shall introduce here novel definitions required for this goal. First we review the formal definitions of kinship and inbreeding coefficients. Then we define a “local” population for every individual, which allows us to distinguish “structural” inbreeding due to the population structure from the “local” inbreeding that applies to individuals with closely-related parents. We then introduce our generalized *F*_ST_ definition as the mean structural inbreeding coefficient, and show that this definition equals the previous *F*_ST_ definition for independent subpopulations. We also generalize previous formulas for changing the reference ancestral population for kinship and inbreeding coefficients. Lastly, we review the connection between kinship coefficients and the covariance of genotypes.

### 3.1 Overview of data and model parameters

Table 1 summarizes the notation used in this work. Our models assume that genotypes at every locus evolve neutrally—by random drift only, in the absence of recent mutation and selection. Thus, only the population structure shapes the covariance structure of genotypes.

**Table 1:** Mathematical notation.

| Symbol | Sec. | Definition |
| --- | --- | --- |
| $x_{ij}$ | 3.1 | Genotype at locus $i$ of individual $j$, counting the number of reference alleles (0,1,2). |
| $p_i^T$ | 3.1 | Reference allele frequency at locus $i$ for population $T$. |
| $f_j^T$ | 3.1 | Inbreeding coefficient: probability that the two alleles at a random locus of individual $j$ are identical by descent (IBD) when the ancestral population is $T$. |
| $\varphi_{jk}^T$ | 3.1 | Kinship coefficient: probability that two alleles drawn randomly one from individual $j$ and the other from individual $k$ at a random locus are IBD when the ancestral population is $T$. |
| $f_S^T$ | 3.1 | Inbreeding coefficient of the panmictic population $S$ when the ancestral population is $T$. |
| $L_j$ | 3.2 | Local population of individual $j$. |
| $L_{jk}$ | 3.2 | Jointly local population of individuals $j$ and $k$ (MRCA population of $L_j$ and $L_k$). |
| $f_j^{L_j}$ | 3.2 | Local inbreeding coefficient of individual $j$ (special case of $f_j^T$ with $T = L_j$). |
| $\varphi_{jk}^{L_{jk}}$ | 3.2 | Local kinship coefficient of individuals $j$ and $k$ (special case of $\varphi_{jk}^T$ with $T = L_{jk}$). |
| $f_{L_j}^T$ | 3.2 | Structural inbreeding coefficient of individual $j$ when the ancestral population is $T$ (special case of $f_S^T$ with $S = L_j$). |
| $f_{L_{jk}}^T$ | 3.2 | Structural kinship coefficient of individuals $j$ and $k$ when the ancestral population is $T$ (special case of $f_S^T$ with $S = L_{jk}$). |
| $F_{ST}$ | 3.3 | Generalized $F_{ST}$: weighted mean $f_{L_j}^T$ over the individuals in a sample. |
| $\bar{t}_T$ | 4 | Mean coalescence time for two alleles at a random locus drawn from $T$. |
| $\bar{t}_j$ | 4 | Mean coalescence time for the two alleles at a random locus of individual $j$. |
| $\bar{t}_{jk}$ | 4 | Mean coalescence time for two alleles drawn at random from each of two individuals $j$ and $k$. |
| $\pi_{ij}$ | 5 | Individual-specific allele frequency (IAF) at locus $i$ of individual $j$. |
| $\theta_{jk}^T$ | 5 | Coancestry coefficient of individuals $j$ and $k$ when the ancestral population is $T$ (equivalent to $\varphi_{jk}^T$ when $j \neq k$ and to $f_j^T$ when $j = k$). |
| $S_u$ | 6 | Supopulation $u$, or intermediate subpopulation $u$ (when ancestral to admixed individuals). |
| $q_{ju}$ | 6 | Admixture proportion of individual $j$ for intermediate subpopulation $S_u$. |
| $\vartheta_{uv}^T$ | 6.2 | Coancestry coefficient of the intermediate subpopulations $S_u$ and $S_v$ when the ancestral population is $T$. |

Let *x*_*ij*_ be observed biallelic genotypes for locus *i* ∈ {1,…, *m*} and diploid individual *j* ∈ {1,…, *n*}. Given a chosen reference allele at each locus, genotypes are encoded as the number of reference alleles: *x*_*ij*_ = 2 is homozygous for the reference allele, *x*_*ij*_ = 0 is homozygous for the alternative allele, and *x*_*ij*_ = 1 is heterozygous. We focus on biallelic loci since they vastly outnumber other types of genetic variants in humans. Note that a multiallelic model, which would require additional notation, could follow in analogy to previous *F*_ST_ work for populations [23].

We assume the existence of a panmictic ancestral population *T* for all individuals under consideration. *T* is generally not required to be the MRCA population, so many choices of *T* are possible. Note that *T* is a collection of organisms ancestral to a given set of individual organisms, shared by all loci, and it is not assumed that the alleles at a given locus coalesce in *T*. Two alleles are said to be “identical by descent” (IBD) if they originate from a single ancestor organism that lived more recently than the given ancestral population *T* [4, 6, 66]. In other words, relationships that precede *T* in time do not count as IBD, while relationships since *T* count toward IBD probabilities. Every locus *i* is assumed to have been polymorphic in *T*, with an ancestral reference allele frequency 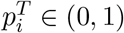, and no new mutations have occurred since then.

The inbreeding coefficient of individual *j* relative to *T*, 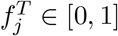, is defined as the probability that the two alleles of any random locus of *j* are IBD when the ancestral population is *T* [67]. Therefore, 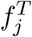 measures the amount of relatedness within an individual, or the extent of dependence between its alleles at each locus. Similarly, the kinship coefficient of individuals *j* and *k* relative to *T*, 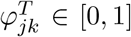, is defined as the probability that two alleles at any random locus, each picked at random from each of the two individuals, are IBD when the ancestral population is *T* [5]. 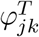 measures the amount of relatedness between individuals, or the extent of dependence across their alleles at each locus. Note that children *j* of parents (*k, l*) have an expected 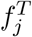 of 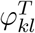 [5]. Both 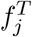 and 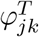 combine relatedness due to the population structure with recent or “local” relatedness, such as that of family members [68]. The values of 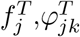 are functions of the chosen ancestral population *T*, which determines the level of relatedness that is treated as unrelated [4, 66]. Thus, 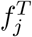 and 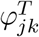 increase if *T* is an earlier rather than a more recent population. The expression “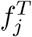 relative to *T*” refers to the value of 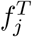 when *T* is chosen as the reference ancestral population [6,66]. The mean 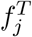 is positive in a structured population [67], and it also increases slowly over time in finite panmictic populations due to genetic drift [69].

Given an ancestral population *T* (not necessarily the MRCA population in this context) and an unstructured subpopulation *S* that evolved from *T*, Malécot defined *F*_ST_ as the mean 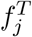 over the individuals in *S* relative to *T* [5], and which we denote by 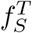. When *S* is itself structured, Wright defined three coefficients that connect *T*, *S* and individuals *I* in *S* [6]: *F*_IT_ (“total inbreeding”) is the mean 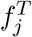 of individuals (*I*) relative to *T*; *F*_IS_ (“local inbreeding”) is the mean 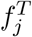 of individuals (*I*) relative to *S*, which Wright did not consider to be part of the population structure; lastly, *F*_ST_ (“structural inbreeding”) is the mean 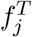 relative to *T* that would result if individuals in *S* mated randomly (and which equals our 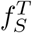). The special case *F*_IS_ = 0 gives *F*_ST_ = *F*_IT_ [6]. See Supplementary Information, Section S1.1 for a more detailed review of these definitions. Wright created the distinction between *F*_ST_ and *F*_IT_ with animal breeding in mind, since mating systems for artificial selection could cause the local inbreeding (*F*_IS_) and therefore also *F*_IT_ to be large at times, but *F*_ST_ measures the more relevant mean inbreeding that results after random mating resumes in the strain [67]. However, in large, natural populations *F*_IS_ is small so *F*_ST_ ≈ *F*_IT_ in these cases. The *F*_ST_ definition has been extended to a set of disjoint subpopulations, where it is the average *F*_ST_ of each subpopulation from the last common ancestral population [23, 24].

In practice, the ancestral population *T* is usually not identified explicitly, which obscures its role in estimating kinship and *F*_ST_. Here we clarify this important matter. Every population of mating organisms can be modeled as descending from a panmictic ancestral population *T*—whether real or a mathematical construct—that at every locus contained the pool of ancestral alleles that modern individuals inherited. By default, the recommended choice of *T* is the MRCA population of the individuals in the sample [22–24, 66, 70]. For example, if all individuals are drawn from one effectively panmictic population, then this population is the MRCA. In a pedigree with unrelated founders, the MRCA population consists of these founders [6, 31]. In a population structure defined by a tree, the MRCA population is the root node at which the first split occurs (Fig. 2). The choice of *T* sets the minimum possible value of 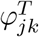: a pair of unrelated individuals drawn from *T* have 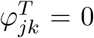, and an individual from *T* (with unrelated parents by definition) has 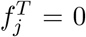 [71]. Thus, assuming that 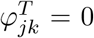 pairs are present in a sample, the set of 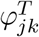 values is in terms of the MRCA population *T* if and only if min 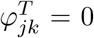. If min 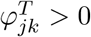, then *T* is more ancestral than the MRCA population. Estimates with min 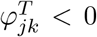—impossible if 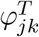 is a probability—have an implicit *T* that is more recent than the MRCA population and cannot be interpreted biologically. For humans, if we ignore the limited Neanderthal and Denisovan introgressions [42, 43], the MRCA population is the real population estimated to have existed in Africa ≈100-200 thousand years ago [32, 33, 40], which first split into the ancestral southern African KhoeSan population (who speak unique “click languages”) and the rest of humans [32, 33, 37, 38, 40].

### 3.2 Local populations

Our generalized *F*_ST_ definition depends on the notion of a local population. Our formulation includes as special cases the independent subpopulations and admixture models, and its generality is in line with recent efforts to model population structure on a fine scale [72, 73], through continuous spatial models [27, 74–76], or in a manner that makes minimal assumptions [56].

We define the *local population L*_*j*_ of an individual *j* as the MRCA population of *j*. In the simplest case, if *j*’s parents belong to the same panmictic subpopulation *S*, then *S* = *L*_*j*_. However, if *j*’s parents belong to different subpopulations, then *L*_*j*_ is modeled as an admixed population (see example below). More broadly, *L*_*j*_ is the most recent panmictic population from which individual *j* drew its alleles and its inbreeding coefficient 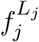 can be meaningfully defined. We define the “local” inbreeding coefficient of *j* to be 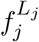, and *j* is said to be *locally outbred* if 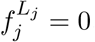.

For any population *T* ancestral to *L*_*j*_, the parameter trio 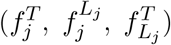 are individual-level analogs of Wright’s trio (*F*_IT_, *F*_IS_, *F*_ST_) defined for a subdivided population [6], with *L*_*j*_ playing the role of *S*. Moreover, just like Wright’s coefficients satisfy

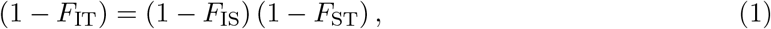

our individual-level parameters satisfy

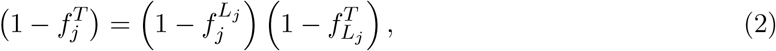

since the probability of the absence of IBD of *j* relative to *T* (which is 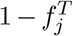)equals the product of the independent probabilities of absence of IBD at two levels: of *j* relative to *L*_*j*_ (which is 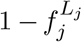), and of *L*_*j*_ relative to *T* (which is 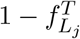). Note that an individual *j* is locally outbred 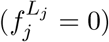 if and only if 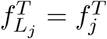.

Similarly, we define the *jointly local population L*_*jk*_ of the pair of individuals *j* and *k* as the MRCA population of *j* and *k*. Hence, *L*_*jk*_ is ancestral to both *L*_*j*_ and *L*_*k*_ (Fig. 2B). We define the “local” kinship coefficient to be 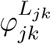, and *j* and *k* are said to be *locally unrelated* if 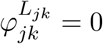. Since the expected inbreeding coefficient of an individual is the kinship of its parents [5], it follows that locally-unrelated parents have locally-outbred offspring.

Consider an individual *j* in an admixture model, deriving alleles from two distinct subpopulations *A* and *B* with proportions *q*_*jA*_ and *q*_*jB*_ = 1 − *q*_*jA*_. Then *L*_*j*_ is modeled as a population that at locus *i* has a reference allele frequency of 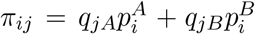, where 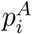 and 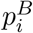 are the allele frequencies in *A* and *B*, respectively. Considering a pair of individuals (*j, k*) and varying their admixture proportions, their jointly local population at one extreme is *L*_*jk*_ = *L*_*j*_ = *L*_*k*_ if and only if *q*_*jA*_ = *q*_*kA*_ (in other words, these individuals have the same local population if and only if their admixture proportions are the same); at the other extreme *L*_*jk*_ is the MRCA population of *A* and *B* if and only if *q*_*jA*_ = 1 and *q*_*kA*_ = 0 or vice versa (in other words, these individuals have the most distant jointly local population if and only if they are not admixed and belong to opposite subpopulations).

### 3.3 The generalized *F*_ST_ for arbitrary population structures

Recall the individual-level analog of Wright’s *F*_ST_ is 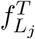, which measures the inbreeding coefficient of individual *j* relative to *T* due exclusively to the population structure (Fig. 2B, Table 1 and Section 3.2). We generalize *F*_ST_ for a set of *n* individuals as

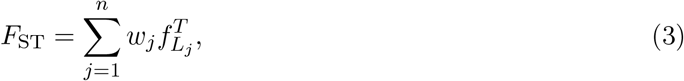

where the most meaningful choice of *T* is the MRCA population of all individuals under consideration, and 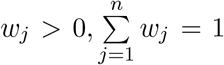 are fixed weights for these individuals. The simplest weights are 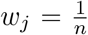 for all *j*. However, we allow for flexibility in the weights so that one may assign them to reflect how individuals were sampled, such as a skewed or uneven sampling scheme. For example, if there are two local populations and the first has twice as many samples as the second, then this can be counteracted by weighing every individual from the first local population half as much as every individual from the second local population. In general, individuals can be weighted inversely proportional to their local population’s sample sizes, a scheme used implicitly in the Hudson pairwise *F*_ST_ estimator [24] and which we iterated for a hierarchy of subdivisions in our analysis of the Human Origins dataset [54]. However, for complex population structures without discrete subpopulations and no obvious sampling biases relative to geography or other variables, we favor uniform weights over complicated weighing schemes (the admixed Hispanic individuals were weighted uniformly in [54]).

This generalized *F*_ST_ definition summarizes the population structure with a single value, intuitively measuring the average distance of every individual from *T*. Moreover, our definition contains the previous *F*_ST_ definition as a special case, as discussed shortly. For simplicity, we kept Wright’s traditional *F*_ST_ notation rather than using something that resembles our 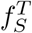 notation. A more consistent notation could be 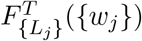, which more clearly denotes the weighted average of 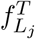 across individuals. Our definition is more general because the traditional *S* population is replaced by a set of local populations {*L*_*j*_}, which may differ for every individual.

#### 3.3.1 Mean heterozygosity in a structured population

Our generalized *F*_ST_ parametrizes the reduction in mean heterozygosity relative to the ancestral population *T* for arbitrary population structures, thus generalizing the familiar connection of the classical *F*_ST_ to allele fixation in an independently-evolving subpopulation. Here we will assume locally-outbred individuals, for which 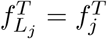. The expected proportion of heterozygotes *H*_*ij*_ of an individual with inbreeding coefficient 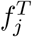 at locus *i* with an ancestral allele frequency 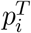 is given by [67]

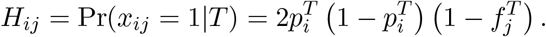

The weighted mean of these expected proportion of heterozygotes across individuals, 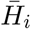, is given by our generalized *F*_ST_:

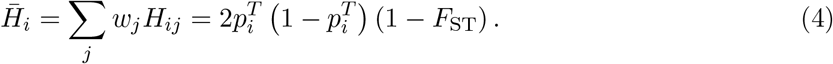

Hence, individuals have Hardy-Weinberg proportions at every locus 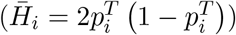 if and only if *F*_ST_ = 0, which in turn happens if and only if 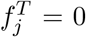 for each *j*. In the other extreme, individuals have fully-fixated alleles at every locus 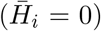, if and only if *F*_ST_ = 1, which in turn happens if and only if 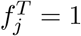 for each *j*.

Eq. (4) presents an apparent paradox since a given sample estimate of the heterozygosity 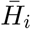 on one side does not depend on *T*, while *F*_ST_ and 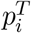 on the other side vary depending on our choice of ancestral population *T*. In fact, both sides of Eq. (4) are constant with respect to *T* under our model: *F*_ST_ increases as *T* is taken to be a more distant ancestral population, but 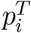 also changes so that 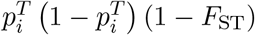 is constant in expectation (see Supplementary Information, Section S4 for a proof of this result).

#### 3.3.2 *F*_ST_ under the independent subpopulations model

Here we show that our generalized *F*_ST_ contains as a special case the currently-used *F*_ST_ definition for independent subpopulations. As discussed above, *F*_ST_ estimators often assume the independent subpopulations model, in which the population is divided into *K* non-overlapping subpopulations that evolved independently from their MRCA population *T* [22–24]. For simplicity, individuals are often further assumed to be locally outbred and locally unrelated. These assumptions result in the following block structure for our parameters,

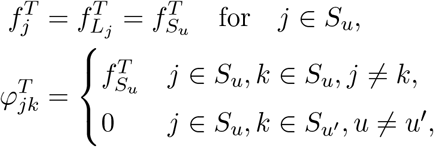

where *j, k* ∈ {1,…, *n*} index individuals, *S*_*u*_, *S*_*u*′_ are disjoint subpopulations treated as sets containing individuals, and *u, u*′ ∈ {1,…, *K*} index these subpopulations. This population structure corresponds to a tree in which every subpopulation split from *T* at the same time (Fig. 2A), which is the required demographic scenario that leads to probabilistically-independent subpopulations.

The generalized *F*_ST_ applied to independent subpopulations agrees with the previous *F*_ST_ definition of the mean per-subpopulation *F*_ST_ [23, 24]:

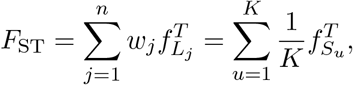

where the weights *w*_*j*_ are such that 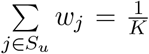. Note also that the *S*_*u*_ for *u* ∈ {1,…, *K*} act as the *K* unique local populations, where *L*_*j*_ = *S*_*u*_ whenever *j* ∈ *S*_*u*_.

### 3.4 IBD probabilities with respect to a reference ancestral population

In developing the generalized *F*_ST_, we have made use of equations that relate IBD probabilities in a hierarchy. Here we generalize these equations to individual inbreeding and kinship coefficients, which allow for transformations of these probabilities under a change of reference ancestral population. Our relationships are straightforward generalizations of Wright’s equation relating *F*_IT_, *F*_IS_, and *F*_ST_ in Eq. (1), now more generally applicable.

Let *A* be a population ancestral to population *B*, which is in turn ancestral to population *C*. The inbreeding coefficients relating every pair of populations in {*A*, *B*, *C*} satisfy

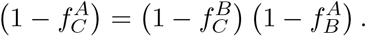

A similar form applies for individual inbreeding and kinship coefficients given relative to populations *A* and *B*, respectively,

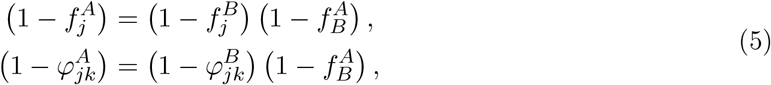

which generalizes Eq. (2). All of these cases follow since the absence of IBD of *C* (or *j*, or *j, k*) relative to *A* requires independent absence of IBD at two levels: of *C* (or *j*, or *j, k*) relative to *B*, and of *B* relative to *A*. All of the above equations can be extended to a multi-level hierarchy just like Wright did for Eq. (1), by iterating at each level [6].

### 3.5 Genotype moments under the kinship model

In the kinship model [5, 6, 67, 77], genotypes *x*_*ij*_ are random variables with first and second moments given by

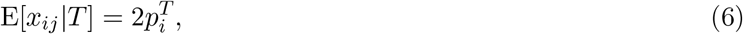

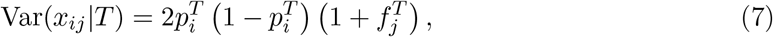

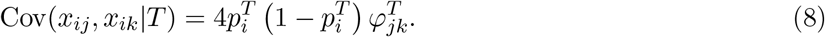

Eq. (6) is a consequence of assuming no selection or new mutations, leaving random drift as the only evolutionary force acting on genotypes [67]. Eq. (7) shows how inbreeding modulates the genotype variance: an outbred individual relative to *T* 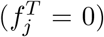 has the Binomial variance of 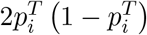 that corresponds to independently-drawn alleles; a fully inbred individual 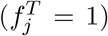 has a scaled Bernoulli variance of 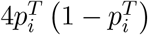 that corresponds to maximally correlated alleles [6]. Lastly, Eq. (8) shows how kinship modulates the correlations between individuals: unrelated individuals relative to *T* 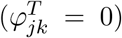 have uncorrelated genotypes, while 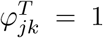 holds for the extreme of identical and fully inbred twins, which have maximally correlated genotypes [5, 77]. Hence, 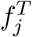 and 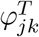 parametrize the frequency of non-independent allele draws within and between individuals. The “self kinship”, arising from comparing Eq. (7) to the *j* = *k* case in Eq. (8), implies 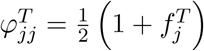, which is a rescaled inbreeding coefficient resulting from comparing an individual with itself or its identical twin.

## 4 Kinship and the generalized *F*_ST_ in terms of the coalescent

Slatkin (1991) [78] derived an expression for the classical *F*_ST_ (for a subdivided population) in terms of mean coalescence times,

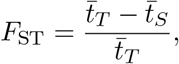

where 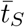 is the mean coalescence time for alleles at a random locus within a subpopulations *S*, and 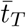 is the mean coalescence time for alleles at a random locus across subpopulations. Here we generalize this expression to encompass inbreeding and kinship coefficients, as well as the generalized *F*_ST_.

In all cases that follow, we generalize 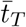 to denote the mean coalescence time for two alleles at a random locus drawn from the ancestral population *T*; in practice it corresponds to the mean coalescence time of the alleles of the two most distant individuals in the sample. The inbreeding and kinship coefficients are given by

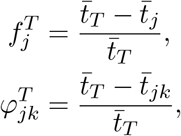

where 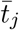 is the mean coalescence time of the two alleles of individual *j* at a random locus, and 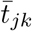 is the mean coalescence time of two alleles drawn at random from each of two individuals *j* and *k* at a random locus (see Supplementary Information, Section S2 for derivations). These mean coalescence times could be estimated as average coalescence times for a large number of neutral loci across the genome. If all individuals in the sample are locally outbred, we obtain the desired expression for the generalized *F*_ST_:

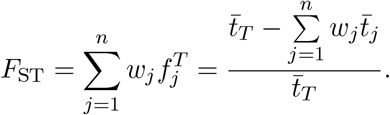

Therefore, the generalized *F*_ST_ equals the relative difference between the weighted mean coalescence times of the alleles within individuals versus the mean coalescence time between the most distantly-related individuals in the sample.

## 5 The coancestry model for individual allele frequencies

*F*_ST_ and its estimators are most often studied in terms of subpopulation allele frequencies [22–24, 57]. Here we introduce a coancestry model for individuals, which is based on *individual-specific allele frequencies* (IAFs) [55, 56] that accomodate arbitrary population-level relationships between individuals. Some authors use the terms “coancestry” and “kinship” exchangeably [23, 70, 71]; in our framework, kinship coefficients are general IBD probabilities (following [68]), and we reserve coancestry coefficients for the IAFs covariance parameters (in analogy to the work of [23]). This coancestry model is the foundation behind the extension of the PSD admixture model we present in Section 6 below, and simplifies the analysis of *F*_ST_ estimator bias in Section 3 of Part II.

In this section we introduce two parameters (see Table 1). First, *π*_*ij*_ ∈ [0, 1] is the IAF of individual *j* at locus *i*. Individual *j* draws its two reference alleles independently with probability *π*_*ij*_. Allowing every locus-individual pair to have a potentially-unique allele frequency allows for arbitrary forms of population structure at the level of allele frequencies [56]. Second, 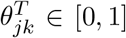 is the coancestry coefficient of individuals *j* and *k* relative to an ancestral population *T*, which modulate the covariance of *π*_*ij*_ and *π*_*ik*_ as shown below.

### 5.1 The coancestry model

In our coancestry model, the IAFs *π*_*ij*_ have the following first and second moments,

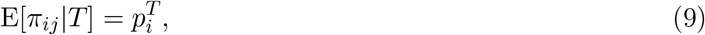

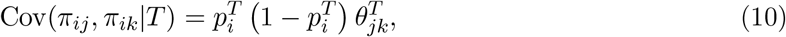

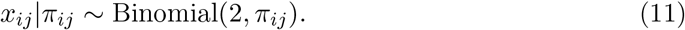

Eq. (9) implies that random drift is the only force acting on the IAFs, and is analogous to Eq. (6) in the kinship model. Eq. (10) is analogous to Eqs. (7) and (8) in the kinship model, with individual coancestry coefficients 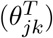 playing the role of the kinship and inbreeding coefficients (for *j* = *k*), a relationship elaborated in the next section. Lastly, Eq. (11) draws the two alleles of a genotype independently from the IAF, which models locally-outbred 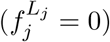 and locally-unrelated 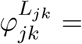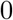 individuals [23]. Hence, the coancestry model excludes local relationships, so it is more restrictive than the kinship model.

Our coancestry model between individuals is closely related to previous models between sub-populations [23, 57]. However, previous models allowed 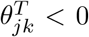[23]. We require that 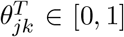 for two reasons: (1) covariance is non-negative in latent structure models [79], such as population structure, and (2) it is necessary in order to relate 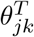 to IBD probabilities as shown next.

### 5.2 Relationship between coancestry and kinship coefficients

Here we show that the coancestry coefficients for IAFs, *θ*_*jk*_, defined above can be written in terms of the kinship and inbreeding coefficients utilized in our more general model. We do so by relating our coancestry coefficients to general kinship coefficients by matching moments. Conditional on the IAFs, genotypes in the coancestry model have a Binomial distribution, so

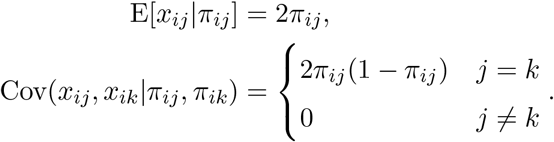

We calculate total moments by marginalizing the IAFs. The total expectation is

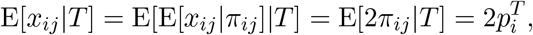

which agrees with Eq. (6) of the kinship model. The total covariance is calculated using

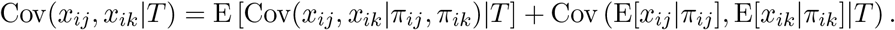

The first term is zero for *j* ≠ *k*, and for *j* = *k* it is

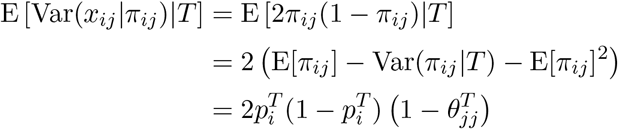

The second term equals 4 Cov (*π*_*ij*_, *π*_*ik*_|*T*) for all (*j*, *k*) cases, which is given by Eq. (10). All together,

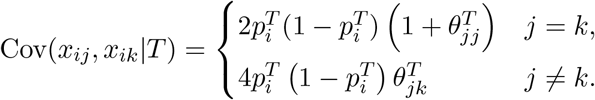

Comparing the above to Eqs. (7) and (8), we find that

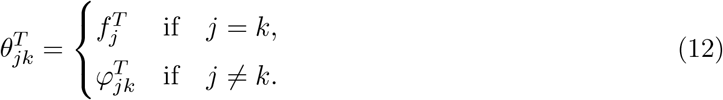

Therefore, our coancestry coefficients are equal to kinship coefficients, except that self-coancestries are equal to inbreeding coefficients.

Since individuals in our IAF coancestry model are locally outbred and locally unrelated, we also have 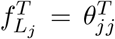 and 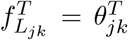 for *j* ≠ *k*. Replacing these quantities in Eq. (3), we obtain the generalized *F*_ST_ in terms of coancestry coefficients.

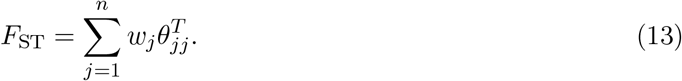

## 6 Coancestry and *F*_ST_ in admixture models

The Pritchard-Stephens-Donnelly (PSD) admixture model [58] is a well-established, tractable model of structure that is more complex than the independent subpopulations model. There are several algorithms available to estimate the PSD model parameters [58–60, 64, 80]. This model assumes the existence of several intermediate ancestral subpopulations, from which individuals draw alleles according to their admixture proportions. However, the PSD model was not developed with *F*_ST_ in mind; we will present a modified model that is compatible with our coancestry model. The results presented in this section are applied to evaluate kinship and *F*_ST_ estimators in Section 6 of Part II, where an admixed population without independent subpopulations is simulated and the true kinship and *F*_ST_ are known.

The PSD model is a special case of our coancestry model with the following additional parameters (see Table 1). The number of intermediate subpopulations is denoted by *K*. Let 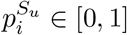 be the reference allele frequency at locus *i* and intermediate subpopulation *S*_*u*_ (*u* ∈ {1,…, *K*}; compare 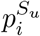 to previous notation 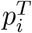 in Table 1). Lastly, *q*_*ju*_ ∈ [0, 1] is the admixture proportion of individual *j* for intermediate subpopulation *S*_*u*_. These proportions satisfy 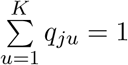 for each *j*.

### 6.1 The PSD model with Balding-Nichols allele frequencies

The original algorithm for fitting the PSD model [58] utilizes prior distributions for intermediate subpopulation allele frequencies and admixture proportions according to

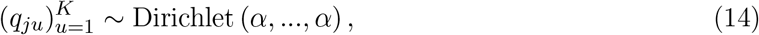

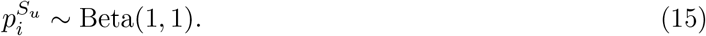

Subsequent work has shown [56, 60] that the PSD model of [58] is then equivalent to forming IAFs

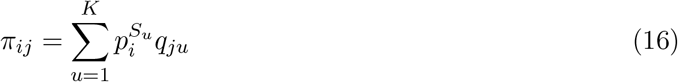

where genotypes are then drawn independently according to *x*_*ij*_|*π*_*ij*_ ~ Binomial(2, *π*_*ij*_).

Here we consider an extension of this model, which we call the “BN-PSD” model, by replacing Eq. (15) with the Balding-Nichols (BN) distribution [7] to generate the allele frequencies 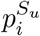 for the intermediate subpopulations from their MRCA population *T*. The BN-PSD model establishes an independent subpopulations structure of the intermediate subpopulations *S*_*u*_ as illustrated in Fig. 4. This combined model has been used to simulate structured genotypes [55, 62, 63], and is the target of some inference algorithms [61, 64]. The BN distribution is the following reparametrized Beta distribution,

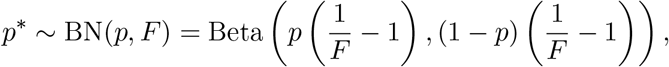

where *p* is the ancestral allele frequency and *F* is the inbreeding coefficient [7]. The resulting allele frequencies *p*^*^ fit into our coancestry model, since E[*p*^*^] = *p* and Var(*p*^*^) = *p*(1 − *p*)*F*.

**Figure 4:**
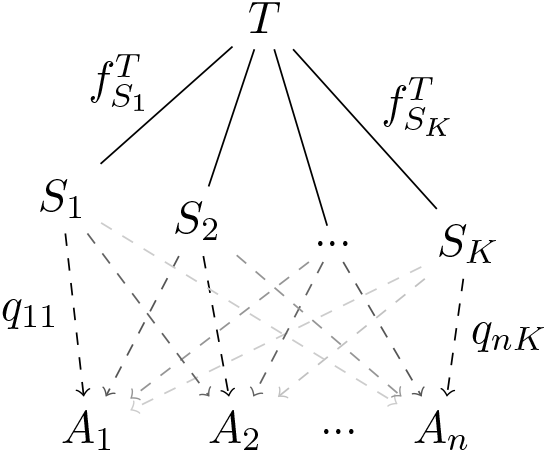
The demographic model of the BN-PSD admixture model. There are *K* intermediate subpopulations *S*_*u*_ for *u* ∈ {1,…, *K*} that evolved independently from their MRCA population *T*, each of which has its own inbreeding coefficient 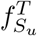 (solid edges). There are *n* admixed individuals denoted as *A*_*j*_ for *j* ∈ {1,…, *n*}, each deriving ancestry from the intermediate subpopulation *S*_*u*_ (dashed arrows with variable shading) in proportion *q*_*ju*_ (i.e. an expected fraction *q*_*ju*_ of alleles of individual *j* are drawn from the intermediate subpopulation *S*_*u*_).

In BN-PSD, the allele frequencies 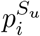 at each locus *i* for intermediate subpopulation *S*_*u*_ are drawn independently from

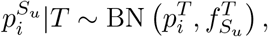

where 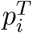 is the ancestral allele frequency and 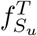 is the inbreeding coefficient of *S*_*u*_ relative to *T* (compare 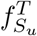 to 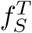 notation in Table 1).

We calculate the coancestry parameters of this model by matching moments conditional on the admixture proportions **Q**= (*q*_*ju*_). We calculate the expectation as

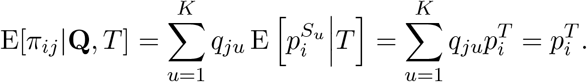

and the IAF covariance is

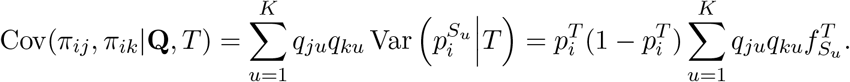

By matching these to Eq. (10), we arrive at coancestry coefficients and *F*_ST_ of

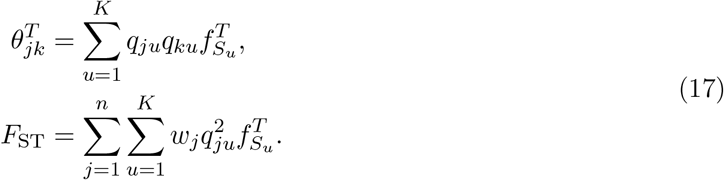

### 6.2 The BN-PSD model with full coancestry

The BN-PSD model contains a restriction that the *K* intermediate subpopulations are independent. Suppose instead that the intermediate subpopulation allele frequencies 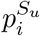 satisfy our more general coancestry model:

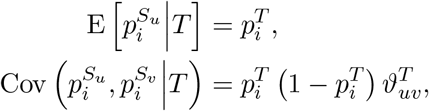

where 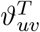 is the coancestry of the intermediate subpopulations *S*_*u*_ and *S*_*v*_. Note that the previous BN-PSD model satisfies 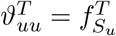 and 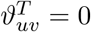 for *u* ≠ *v*. Repeating our calculations assuming our full coancestry setting, individual coancestry coefficients and *F*_ST_ are given by

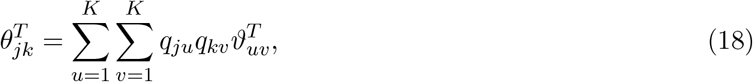

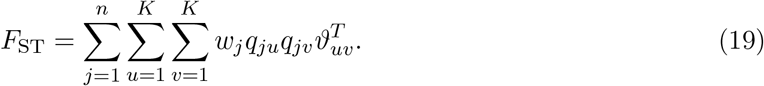

Therefore, all coancestry coefficients of the intermediate subpopulations influence the individual coancestry coefficients and the overall *F*_ST_. The form for 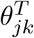 above has a simple probabilistic interpretation: the probability of IBD at random loci between individuals *j* and *k* corresponds to the sum for each pair of subpopulations *u* and *v* of the probability of the pairing (*q*_*ju*_*q*_*kv*_) times the probability of IBD between these subpopulations 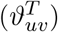. Note that Eq. (18) was derived independently for a related model [81], but the value of *F*_ST_ for a set of admixed individuals—which we provide in Eq. (19)—had not been described before to the best of our knowledge.

## 7 Discussion

We presented a generalized *F*_ST_ definition corresponding to a weighted mean of individual-specific inbreeding coefficients. Compared to previous *F*_ST_ definitions, ours is applicable to arbitrary population structures, and in particular does not require the existence of non-overlapping subpopulations.

We considered two closely-related population structure models with individual-level resolution: the kinship model for genotypes, and our new coancestry model for IAFs (individual-specific allele frequencies). The kinship model is the most general, applicable to the genotypes in arbitrary sets of individuals. Our IAF model requires a local form of Hardy-Weinberg equilibrium, and it does not model locally-related or locally-inbred individuals. Nevertheless, IAFs arise in many applications, including admixture models [59], estimation of local kinship [55], genome-wide association studies [82], and the logistic factor analysis [56]. We prove that kinship coefficients, which control genotype covariance, also control IAF covariance under our coancestry model.

We also calculated *F*_ST_ for admixture models. To achieve this, we framed the PSD (Pritchard-Stephens-Donnelly) admixture model as a special case of our IAF coancestry model, and studied extensions where the intermediate subpopulations are more structured. *F*_ST_ was previously studied in an admixture model under Nei’s *F*_ST_ definition for one locus, where *F*_ST_ in the admixed population is given by a ratio involving admixture proportions and intermediate subpopulation allele frequencies [52]. On the other hand, our *F*_ST_ is an IBD probability shared by all loci and independent of allele frequencies. Under our framework, the *F*_ST_ of an admixed individual is a sum of products, which is quadratic in the admixture proportions and linear in the coancestry coefficients of the intermediate subpopulations. In the future, inference algorithms for our admixture model with fully-correlated intermediate subpopulations could yield improved results, including coancestry and *F*_ST_ estimates.

Our probabilistic model reconnects *F*_ST_ [21, 23, 24] to inbreeding and kinship coefficients [68, 70, 83, 84], all quantities of great interest in population genetics, but which are currently studied in isolation. The main reason for this isolation is that *F*_ST_ estimation assumes the independent sub-populations model, in which kinship coefficients are uninteresting. However, study of the generalized *F*_ST_ in arbitrary population structures requires the consideration of arbitrary kinship coefficients [68]. Our work lays the foundation necessary to study estimation of the generalized *F*_ST_, which is the focus of our next publication in this series (Part II).

## Acknowledgments

This research was supported in part by NIH grant R01 HG006448.

## Supplementary Information

### S1 Review of previous *F*_ST_ definitions

Here we review how *F*_ST_ measures the population structure of individuals, parametrizes genetic drift, and has been adapted to studying the genetic diversity at individual loci. Studies with different goals have demanded various definitions of *F*_ST_ as starting points, which have generated considerable confusion [22, 24, 48, 51, 53, 85]. Here we group these working definitions of *F*_ST_ into three classes and discuss their connection to our work. Note that all previous *F*_ST_ definitions are fundamentally about a subpopulation or a collection of disjoint subpopulations, and do not apply to individuals with arbitrary relatedness such as Hispanics (Section 2 and [54]) who have individual-specific admixture proportions (this admixture model is studied in Section 6 above and utilized to simulate data to benchmark generalized *F*_ST_ estimation in Section 6 of Part II).

#### S1.1 *F*_ST_ as a function of inbreeding coefficients

The original *F*_ST_ of Malécot and Wright is the mean inbreeding coefficient in a subpopulation *S* relative to an ancestral population *T* [5, 6], which corresponds to our 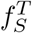. If *S* is unstructured [5], then 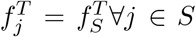 in our notation and thus it can be given in terms of individual inbreeding coefficients by

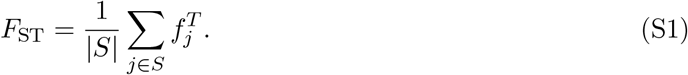

When *S* is structured [6] then three quantities are specified which may be given in terms of individual coefficients by

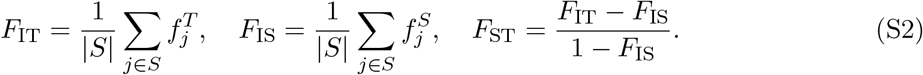

Note that when *S* is unstructured then Eq. (S2) reduces to Eq. (S1), since 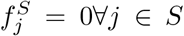, so *F*_IT_ = 0 and *F*_ST_ = *F*_IT_. Additionally, Eq. (S2) holds under our generalized framework since 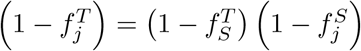, but the alternative form

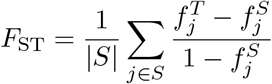

is more directly comparable to our generalized definition of Eq. (3).

This original *F*_ST_ and the earlier inbreeding [67] and kinship coefficients [5] were all estimated from pedigrees rather than genetic markers as it is now more common. Thus, this *F*_ST_ measures only the relatedness of individuals in a subpopulation, it is independent of mutation rates or selection, and it is not defined by any particular genetic marker. Inbreeding coefficients and *F*_ST_ were estimated from a pedigree using the method of path coefficients [3]. Our generalized *F*_ST_—defined in Eq. (3) using individual inbreeding coefficients—corresponds most closely to this original *F*_ST_ definition, with the important exception that we aim to estimate realized kinship coefficients rather than their expected values under the pedigree [86].

#### S1.2 *F*_ST_ as a model parameter of allele variance

Consider a biallelic locus *i* and some allele taken as a reference, which had an allele frequency 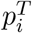 in the ancestral population *T* and which evolves to have an allele frequency 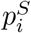 in a subpopulation *S* that derives from *T* such that the mean inbreeding of every individual in *S* relative to *T* is *F*_ST_. *T* and *S* are implicitly panmictic populations, so that genotypes are in Hardy-Weinberg equilibrium and 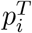 and 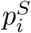 suffice to describe their allele distributions. Wright found that for neutral loci *i* (without mutation and selection) the variance of the possible random values 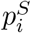 that could result given fixed 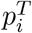 and *F*_ST_ parameters is [31]

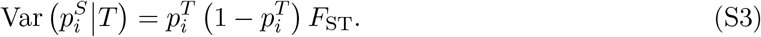

Note again that this equation results from considering the effect that relatedness of individuals has on their allele frequencies at neutral loci, and that *F*_ST_ is thus shared across all such neutral loci.

Many subsequent works have taken Eq. (S3), restated as

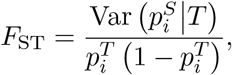

to define *F*_ST_ [19, 22–24, 49, 51, 71, 87]. This alternative definition can lead to confusion for three reasons: it can be mistakenly interpreted as applying to all loci (including loci under mutation or selection); it suggests that every locus *i* has its own *F*_ST_; and it depends on the unknowns 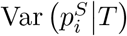 and 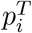 that have been interpreted in various ways [22, 24, 48, 49, 51–53, 88, 89]. We stress that Wright and Malécot originally defined *F*_ST_ from inbreeding coefficients and Eq. (S3) was derived as a consequence of the relatedness of individuals (as captured by *F*_ST_) and applies only to neutral loci [5, 31].

Complicating matters, in developing the effect of *F*_ST_ on allele distributions, both Malécot and Wright extended *F*_ST_ to incorporate the effect of mutation into Eq. (S3), resulting in formulas for the *F*_ST_ in a population at equilibrium, such as

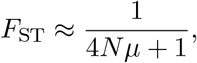

for a population with *N* individuals at all times, where *μ* is the sum of migration (proportion of individuals *N* per generation) and mutation rates (proportion of mutations per locus per generation), and *F*_ST_ above is the approximate value approached in infinite time [5, 6]. Both authors note that migration reduces *F*_ST_ in the inbreeding definition, and mutation has an identical mathematical effect on reducing the variance of 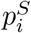, thus mutation reduces the *effective F*_ST_ by reducing the probability of allele fixation [5, 6]. In contrast, the inbreeding *F*_ST_ approaches 1 with infinite time in a finite and isolated population [69], regardless of mutation. Since locus mutation does not alter inbreeding values, this extended *F*_ST_ that captures mutation is no longer compatible with the inbreeding *F*_ST_. Many later works with greater focus on the evolution of allele frequencies than on relatedness adopted the *F*_ST_ definition that incorporates mutation [16, 17, 51, 78, 90–94]. Frameworks that assume neutral loci only—and thus are compatible with the inbreeding *F*_ST_— include our work, method-of-moments *F*_ST_ estimators [17, 22–24, 46, 47, 95] and Normal [23, 57, 96] and Bayesian likelihood models based on the Beta (for biallelic loci) or Dirichlet (multiallelic) distributions [7, 20, 36, 61] for the subpopulation allele frequencies 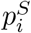. Some authors model *F*_ST_ and mutation as separate effects [97, 98].

In the coalescent framework, in the limit of small mutation rates, *F*_ST_ was shown to equal

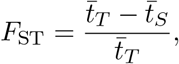

where 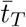 and 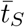 are average coalescence times for alleles within the populations *T* and *S*, respectively [78]. This connection to coalescent times led to the *R*_ST_ statistic that remarkably estimates *F*_ST_ from microsatellites while excluding the effect t of the relatively high mutation rate of these variants under some assumptions [98]. The Weir-Cockerham *F*_ST_ estimator and *R*_ST_ are special cases of *φ*_ST_ in the AMOVA framework [99, 100].

#### S1.3 *F*_ST_ as a data-dependent statistic that measures variance at a locus

Locus-specific *F*_ST_ estimates are often employed to identify loci under selection [14–21]. In this setting, *F*_ST_ is often defined by the following sample estimate of Eq. (S3) per locus *i*,

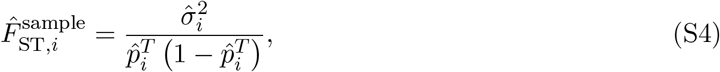

where 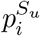 is the reference allele frequency in each subpopulation *S*_*u*_, 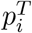 is estimated by the sample mean over the *K* subpopulations 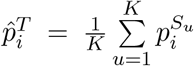, and Var (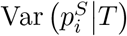 1*T*) is estimated by the sample variance 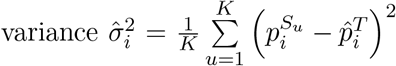 [48, 51–53, 85, 88, 89, 101–106]. Unlike the random variable 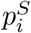 in the *F*_ST_ definition of Eq. (S3), studies of Eq. (S4) usually treat 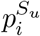 as fixed parameters. Thus, although 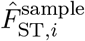 shares many of the properties of *F*_ST_, 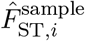 is a biased estimator of the *F*_ST_ from Eq. (S3) [22, 71], so these definitions are not compatible. Nevertheless, since 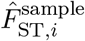 is effective for studying the evolution of individual loci, it has spawned its own field of research, starting with Nei’s *G*_ST_ that generalizes 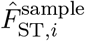 to multiple alleles [48] and is often treated as *F*_ST_ [52, 53, 78, 85, 88, 89, 102, 103, 106], related single-locus *F*_ST_ estimators based on method-of-moments [16–19] or Bayesian models [20, 21, 96, 106], and alternative quantities such as 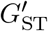 [49] and *D* [50] *G*_ST_ approximates *F*_ST_ better than 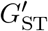 and *D*, especially under a low mutation rate [51]). Note that although *F*_ST_ was previously studied for biallelic loci only [5, 6, 31, 48], there are more recent *F*_ST_ models that generalize Eq. (S3) for neutral multiallelic loci [7, 22, 23] analogous to how the *G*_ST_ statistic generalizes Eq. (S4). Locus-specific *F*_ST_ estimates present unique challenges since their sampling distribution depends on demography and heterozygosity or the maximum allele frequency at the locus [16–20, 49, 53, 104, 107]. The focus of our work is to generalize and accurately estimate the genome-wide *F*_ST_ in individuals with arbitrary relatedness, and does not presently concern locus-specific *F*_ST_ estimation or the identification of loci under selection.

### S2 Derivation of kinship and *F*_ST_ in terms of mean coalescence times

We shall consider the probability of identity by descent (IBD) in a random process that admits mutations along the coalescent tree. Interestingly, the limit as the mutation rate goes to zero results in non-trivial connections between the IBD coefficients and coalescence times. Our proof closely mirrors that of [78].

Let *μ* be the mutation rate, in units of mutations per base per generation, which is assumed to be a constant for all branches of the tree. Let *h*_1_ and *h*_2_ denote two haploid DNA sequences (we shall convert to diploid individuals in the end). Let 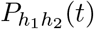 be the probability that *h*_1_ and *h*_2_ coalesce in generation *t*. By definition, the sum of these probabilities across all coalescence times (*t* ≥ 1) equals one:

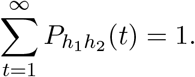

The overall probability that a given random locus at both sequences is IBD is the expectation of (1 − *μ*)^2*t*^—the probability that a mutation has not occured by generation *t* at this locus for both sequences *h*_1_ and *h*_2_:

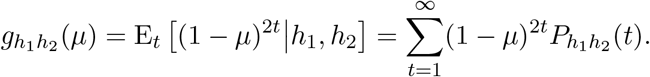

Note that 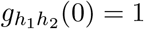 and

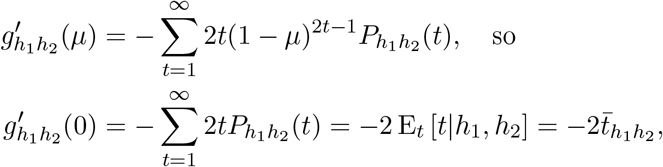

where 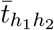 is the mean coalescence time of sequences *h*_1_ and *h*_2_. To proceed, consider the equivalent quantity for the two most distant sequences in the sample, which are taken as being drawn independently from the ancestral population *T*:

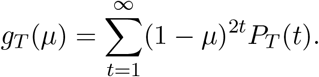

The IBD coefficient of interest, 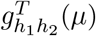, is a relative probability related to 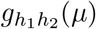 and *g*_*T*_ (*μ*) in the same manner as *F*_ST_, *F*_IT_ and *F*_IS_, namely

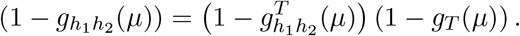

Note that solving for 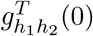 above gives an undefined value (0/0), since 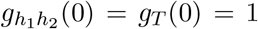. Nevertheless, solving for 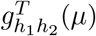 first (for *μ* ≠ 0) and taking the limit as the mutation rate goes to zero (using L’Hôpital’s rule), we obtain the IBD probability of interest, for *h*_1_ and *h*_2_ relative to *T*:

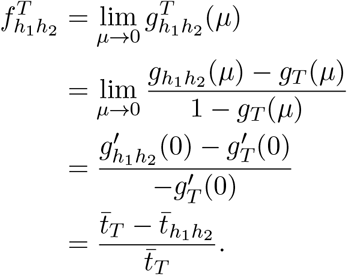

The coefficients of interest are special cases of the last expression, as follows. The inbreeding coefficient is

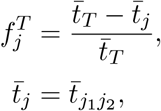

where *j*_1_ and *j*_2_ are the two haplotypes of individual *j* (the maternal and paternal alleles). Similarly, the kinship coefficient is an average of haplotype comparisons across individuals,

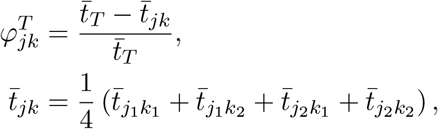

where *j*_1_ and *j*_2_ are the two haplotypes of individual *j*, and *k*_1_ and *k*_2_ are the two haplotypes of individual *k*.

### S3 Empirical Bayes estimation of subpopulation allele frequencies for map

The allele frequencies shown in the map of Fig. 1B are estimated from genotypes using Empirical Bayes with a Beta prior [108], as follows. Let *x*_*ij*_ be the number of reference alleles at locus *i* and subpopulation *j*, and *n*_*ij*_ be the total number of alleles. We model the desired subpopulation allele frequencies *π*_*ij*_ as drawn independently from a Beta prior:

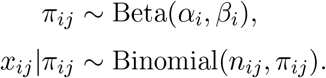

The marginal distribution of *x*_*ij*_ is the Beta-Binomial. The posterior estimate of *π*_*ij*_ that was displayed in Fig. 1B is

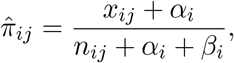

which compared to the sample estimate 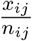 is “shrunk” toward the prior mean 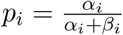 depending on sample size (*n*_*ij*_ ≫ *α*_*i*_ + *β*_*i*_ have 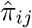 close to 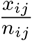, while *n*_*ij*_ ≪ *α*_*i*_ + *β*_*i*_ have 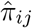 closer to *p*_*i*_).

Instead of choosing *α*_*i*_, *β*_*i*_ a priori, in empirical Bayes estimation *α*_*i*_, *β*_*i*_ are the values that maximize the log-likelihood of the data,

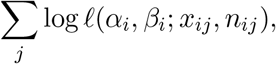

where *ℓ* is the Beta-Binomial likelihood function.

The Human Origins dataset was processed as described in [54], and additionally filtered to consider only loci with a minor allele frequency ≥ 10% (362,437 loci) in identifying the locus with the median per-locus Weir-Cockerham *F*_ST_ estimate (using the *K* = 244 sub-subpopulations to partition individuals). For the locus rs2650044 displayed on Fig. 1B we estimated *α*_*i*_ ≈ 1.83 and *β*_*i*_ ≈ 8.34.

### S4 Proof that expected heterozygosity is indepenent of *T*

Here we show that *H*_*ij*_ has the same form conditioned on some ancestral population *S* as it does for any other choice *T* ancestral to *S*. Conditional on *T*, all of 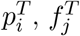 and 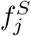 are constant parameters, but 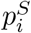 is a random allele frequency that drifted from the more ancestral 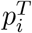 frequency, so 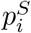 must be marginalized. Therefore, it suffices to prove that

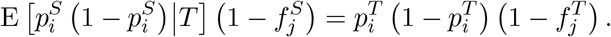

We assume that 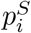 satisfies the coancestry model of Eqs. (9) and (10), yielding:

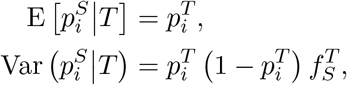

where 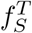 is the inbreeding coefficient of population *S* relative to *T* (see Section 3.1) and satisfies the IBD shift identity in Eq. (5):

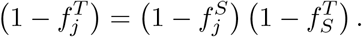

The desired conclusion follows:

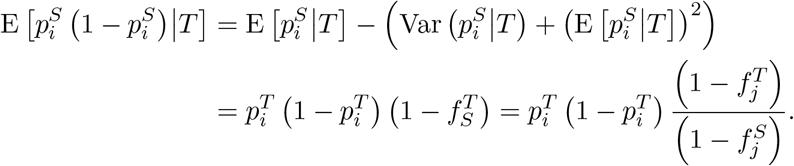

